# Metabolic evidence for distinct pyruvate pools inside plant mitochondria

**DOI:** 10.1101/2021.10.30.466557

**Authors:** Xuyen H. Le, Chun-Pong Lee, Dario Monachello, A. Harvey Millar

**Author notes:** **Author Contributions:** X.H.L., C.P.L., and A.H.M. designed the research. X.H.L. performed most of the experiments and data analysis, C.P.L. assisted with some of the mass spectrometry and data analysis, D.M. performed the interactome analyses. X.H.L., C.P.L., and A.H.M. wrote the paper.

## Abstract

The majority of the pyruvate inside plant mitochondria is either transported into the matrix from the cytosol via the mitochondria pyruvate carrier (MPC) or synthesised in the matrix by alanine aminotransferase (AlaAT) or NAD-malic enzyme (NAD-ME). Pyruvate from these origins could mix into a single pool in the matrix and contribute indistinguishably to respiration, or they could maintain a degree of independence in metabolic regulation. Here, we demonstrated that feeding isolated mitochondria with U-^13^C-pyruvate and unlabelled malate enables the assessment of pyruvate contribution from different sources to TCA cycle intermediate production. Imported pyruvate is the preferred source for citrate production even when the synthesis of NAD-ME-derived pyruvate was optimised. Genetic or pharmacological elimination of MPC activity removed this preference and allowed an equivalent amount of citrate to be generated from the pyruvate produced by NAD-ME. Increasing mitochondrial pyruvate pool size by exogenous addition only affected metabolites from pyruvate transported by MPC whereas depleting pyruvate pool size by transamination to alanine only affected metabolic products derived from NAD-ME. Together, these data reveal respiratory substrate supply in plants involves distinct pyruvate pools inside the matrix that can be flexibly mixed based on the rate of pyruvate transport from the cytosol. These pools are independently regulated and contribute differentially to organic acids export from plant mitochondria.

**Significance statement:** Pyruvate is the primary respiratory substrate for energy production to support plant growth and development. However, it is also the starting material of many other pathways. Prioritisation of respiratory use over other competing pathways would enable a level of control when pyruvate is delivered to mitochondria via the mitochondrial pyruvate transporter. We demonstrated the existence of two distinct pyruvate pools in plant mitochondria suggesting inner mitochondrial organisation allows metabolic heterogeneity, hence metabolic specialisation. This explains why NAD-ME flux into plant respiration is low and confirms the prominent link between imported pyruvate and energy production. This compartmentation also reveals how NAD-ME supplies substrate to the mitochondrial pyruvate exporter in plants, especially during C4 metabolism.

## Introduction

Pyruvate is the main product of cytosolic glycolysis and the fuel for aerobic respiration in most organisms (1). It is oxidised by the mitochondrial pyruvate dehydrogenase complex (PDC), producing acetyl-CoA which enters the tricarboxylic acid (TCA) cycle and generates reductant equivalent for ATP production. In plants, the majority of pyruvate in the mitochondrial matrix is supplied from three sources: (i) transport from the cytosol via the mitochondria pyruvate carrier (MPC), (ii) alanine aminotransferase (AlaAT) that interconverts alanine and pyruvate, and (iii) oxidative decarboxylation of malate via NAD-malic enzyme (NAD-ME). We have recently shown that the combined action of MPC and the AlaAT is responsible for providing the bulk of pyruvate to the TCA cycle in *Arabidopsis thaliana*, while NAD-ME only has a minor role under both *in vivo* and *in vitro* conditions (2). This observation is in contrast to the long-standing view that NAD-ME is a major contributor to pyruvate-dependent, TCA-cycle linked respiration due to high malate availability and oxidation rate compared to other respiratory substrates and the low level of pyruvate in plant cells (3–5). But is consistent with *in vivo* labelling studies has revealed ME activity accounts for only 3% of the pyruvate synthesized in respiring maize root tips (6, 7) and 1% in *Xanthium strumarium* leaves (8). It is plausible that views about the centrality of the respiratory role of malic enzyme have been significantly influenced by its activity at low pH in purified enzyme samples or isolated mitochondria (7).

We found mitochondria isolated from an MPC1 loss-of-function mutant (*mpc1*) possess a higher NAD-ME-dependent pyruvate production rate (2), indicating that a metabolic switch may exist to control pyruvate supply and usage in order to meet specific metabolic and energy demands under different conditions. One possible explanation for this phenomenon is that plant mitochondria operate separate pyruvate pools: an imported pyruvate pool that sustains the TCA cycle, and NAD-ME-derived pyruvate that serves as an emergency valve and is only switched into TCA cycle metabolism when pyruvate import into the matrix is insufficient to satisfy cellular energy demand.

One known mechanism in plant cells to separate metabolic pools is substrate channelling, either (i) direct channel formed by interacting enzymes or (ii) increasing the local concentration of enzymes by bringing enzymes together into clusters, rather than having them distributed through the cell. It preferably facilitates the transfer of substrates from one active site of one enzyme to the other sequential enzymes with minimal mixing substrates to the common pool to minimize usage of substrates by competed pathways (9). The occurrence of substrate channelling has been studied in the TCA cycle, making it one of the preceded examples of metabolic direct sequential channelling (10, 11). The associative pairing of sequential enzymes, called a metabolon has a kinetic advantage over free enzymes scouting for substrates from a single metabolic pool (12), for example between malate dehydrogenase (MDH), citrate synthase (CS) (12–16). Similarly, purinosome that is responsible for *de novo* purine synthesis is an enzyme cluster in which reaction rate is enhanced by increased enzyme concentration is probabilistic rather than direct (17). It was also suggested by the clustering model that flux would increase by 6-fold for a 2-step pathway and over 100-fold for a 3-step pathway compared to freely diffusing enzymes within a cell (18).

In this study, we explored the evidence for channelling-like phenomena in respiratory metabolism before the TCA cycle, in the import of pyruvate via MPC and its delivery to the TCA cycle. This necessitated the use of an intact mitochondrial system to allow pyruvate transport and use by respiratory metabolism. We assessed the case for channelling by monitoring how imported pyruvate interacts and competes with pyruvate generated by NAD-ME in the matrix using co-feeding of mitochondria with multiple labelled and unlabelled substrates. Our results revealed a preferential usage of transported pyruvate rather than NAD-ME-derived pyruvate by pyruvate dehydrogenase complex (PDC) and TCA cycle enzymes which could be reversed when MPC1 was absent or its activity chemically inhibited. This flexibility indicates the presence of separate pyruvate pools in the matrix and suggests the occurrence of a regulatory system that is more flexible than physical substrate channelling of pyruvate between MPC and PDC, more akin to the proposed compartmentation of mitochondrial metabolism and the apparent movement of metabolites between compartments in models of mammalian cell metabolism (19–21).

## Results

### Transported pyruvate is converted to citrate but when generated from NAD-ME it is preferentially exported from isolated mitochondria

Using selective reaction monitoring-mass spectrometry (SRM-MS) assays to trace the fate of two substrates simultaneously (2, 22), we assessed the relative contribution of MPC1 and NAD-ME to metabolites derived from the pyruvate pool in mitochondria from Col-0. Isolated mitochondria were subjected to a series of ^13^C_3_-pyruvate concentrations ranging from 0 to 500 μM with a fixed concentration of 500 μM malate at pH 6.4 to determine if a high NAD-ME activity competes with MPC for supplying pyruvate to the TCA cycle *in vitro*. This level of acidity maximises NAD-ME activity (23). Under these conditions, imported malate can either be oxidised to oxaloacetate by MDH or oxidised to pyruvate via NAD-ME. Citrate is then synthesised by combining oxaloacetate with acetyl-CoA made either from exogenously supplied pyruvate or from pyruvate formed in the matrix via NAD-ME (24). The pyruvate used in this process can be distinguished by isotopic forms (Figure 1A, 1E). The relative amount of labelled and unlabelled citrate exported to the extra-mitochondrial medium can be used to assess the amount of respiratory pyruvate supplied by MPC and NAD-ME, respectively.

**Figure 1.**
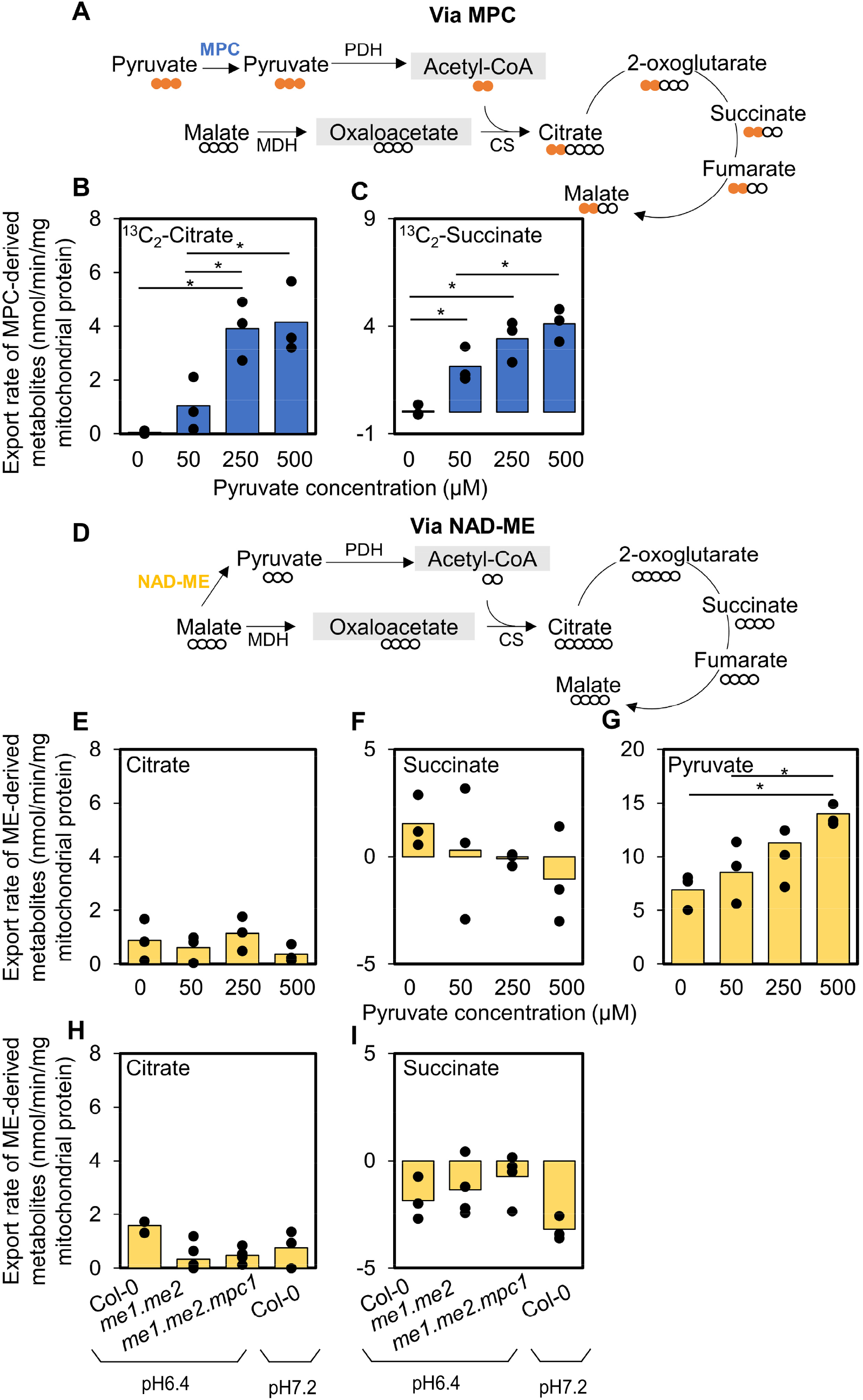
The usage of imported pyruvate for TCA cycle is preferred to that of NAD-ME-derived pyruvate by Col-0. (A-G) Mitochondria were incubated and 0, 50, 250 and 500 μM of ^13^C_3_-pyruvate in the presence of 500 μM malate and ADP at pH 6.4 to increase ME activity. The isotopic incorporation patterns of labelled pyruvate and unlabelled malate into citrate via MPC (A) and via NAD-ME (D) are shown. Bar graphs show export rates of (B) ^13^C_2_-citrate (via MPC), (C) ^13^C_2_-succinate (via MPC); (E) citrate (via NAD-ME), (F) succinate (via NAD-ME), (G) pyruvate (via NAD-ME) of Col-0 mitochondria. (H-I) Col-0, *me1*.*me2* and *me1*.*me2*.*mpc1* mitochondria were incubated in a mixture of 500μM malate and 500μM ^13^C_3_-pyruvate at pH6.4 and pH7.2. Bar graphs compare the export rates of (H) unlabeled citrate (via NAD-ME), (I) unlabeled succinate (via NAD-ME) of Col-0 versus mutant mitochondria. Quantification was carried out using SRM-MS to directly assess substrate consumption and product generation of substrate-fed mitochondria after separating mitochondria from the extra-mitochondrial space by centrifugation through a single silicon oil layer. The rates were calculated from time course values of metabolite concentration recorded in the extra-mitochondrial space after varying incubation periods. Each bar represents averaged value from three or more replicates represented by data points. Significant differences between different pyruvate concentrations and between wildtype and mutants are denoted by asterisks based on Student’s t-tests (*, p < 0.05). Abbreviations: PDH - Pyruvate dehydrogenase, MDH – Malate dehydrogenase, CS – Citrate synthase.

As expected, increasing exogenous ^13^C_3_-pyruvate concentration correlated with enhanced production and export of ^13^C_2_-citrate and ^13^C_2_-succinate due to increased pyruvate import (Figure 1B, 1C). Interestingly, unlabelled pyruvate (via NAD-ME) was released from the mitochondria at high rates (at 14 nmol/min/mg protein with 500 μM provided pyruvate), at least seven fold compared to other unlabelled metabolites (Figure 1G). When a smaller concentration of ^13^C_3_-pyruvate (0 or 50 μM) was supplied, a lower amount of NAD-ME-derived pyruvate from malate exported was detected (7-9 nmol/min/mg protein). However, the amount of unlabelled citrate and succinate exported by mitochondria was independent of the unlabelled pyruvate available (Figure 1E, 1F, Supplemental Figure S1A). From these data, we concluded that the amount of exogenous ^13^C_3_-pyruvate had little impact on the entry of NAD-ME-derived pyruvate into the TCA cycle, suggesting a mechanism to select the origin of pyruvate for entry to the TCA cycle exists in plant mitochondria. Especially when providing mitochondria with 500 μM pyruvate, MPC-derived citrate was more than 4 times higher in concentration than NAD-ME-derived citrate.

In order to confirm the limited entry of NAD-ME-derived pyruvate into the TCA cycle, NAD-ME double mutants *nad*.*me1/nad*.*me2* (*me1*.*me2)* and triple mutants of NAD-ME and the MPC complex *nad*.*me1/nad*.*me2/mpc1* (*me1*.*me2*.*mpc1)* were fed with a mixture of 500 μM pyruvate and 500 μM malate. Our previous study showed that mitochondria from wildtype plants (Col-0) at pH 6.4 showed a rapid increase in unlabelled pyruvate generated from malate by NAD-ME, while *me1*.*me2* and *me1*.*me2*.*mpc1* did not due to the absence of the NAD-ME enzyme and wildtype mitochondria at pH 7.2 also failed to accumulate unlabelled pyruvate (NAD-ME activity is minor) (2). Labelled citrate and downstream labelled TCA metabolites derived from ^13^C_3_-pyruvate imported via MPC were steadily produced and exported, proving that malate oxidation to OAA by MDH, pyruvate oxidation by PDC and other TCA cycle machineries were not defective in these mitochondria (2). Despite this, the rate of unlabelled citrate exported by wildtype at pH 6.4 was extremely low and did not show a significant difference to that by *me1*.*me2* and *me1*.*me2*.*mpc1* and Col-0 at pH 7.2 (Figure 1H), suggesting that pyruvate produced by NAD-ME at pH 6.4 in the matrix was mostly exported and not oxidised by PDC for citrate synthesis (Figure 1E). Other downstream unlabelled metabolites, such as 2-oxoglutarate and succinate, showed similar trends with no significant amounts produced (Figure 1I, Supplemental Figure S1B). This confirms the hypothesis that there is a preference of using imported pyruvate for TCA cycle metabolism and we cannot treat pyruvate from different origins as single pool.

### No evidence for physical interaction of MPC and PDC

To investigate the possibility of substrate channelling which could help explain the observed metabolic compartmentation phenomena, we first looked at reports of protein-protein interaction in the mitochondria. But we found no evidence for PDC subunit proteins binding to MPC1 in plants (25, 26), yeast (27, 28) or mammalian cells (15, 29–32). To look instead for evidence of MPC1 binding to PDC subunits, we conducted a high-throughput yeast two-hybrid (Y2H) assay using MPC1 as bait to screen for interactions with about 12000 proteins in the Arabidopsis library to probe the possible interaction between MPC and PDC (Supplemental Table 1). The library includes the PDC components E1α, E1β, E2 and E3 (At1g59900, At1g24180, At5g50850, At3g52200, At3g13930, At1g54220, At1g48030, At3g17240) and other TCA cycle enzymes. High throughput screening failed to identify any physical interactions of MPC1 with PDC subunits, suggesting the presence of distinct pyruvate pools is not likely to result from a clear physical association that promotes substrate channelling. GRXS17 (At4g04950) and GRX480 (At1g28480) were included as positive control baits in the same screen to validate the experiment, and known interactors were confirmed by colony sequencing (for GRXS17, known interactors BolA2-AT5G09830, BolA4-AT5G17560 and Dre2-AT5G18400 (33); for GRX480, known interactors TGA3-AT1G22070 and TGA7-AT1G77920 (34)).

### The use of mitochondrial NAD-ME-derived pyruvate by PDC is stimulated by eliminating MPC1 activity either genetically or chemically

We next examined if the preference of PDC for MPC-pyruvate over NAD-ME-pyruvate is constitutive or if it changes depending on the pyruvate-supply in isolated mitochondria. We have recently shown that MPC1 is required for pyruvate import from the cytosol into mitochondrial matrix by monitoring the increase in labelled citrate and export of downstream TCA cycle metabolites at pH 7.2 (2). Under conditions that optimise NAD-ME activity (pH 6.4), the loss of MPC1 resulted in the lack of mitochondrial ^13^C-pyruvate import, leading to a substantial reduction in the export of labelled citrate, succinate and malate to the extramitochondrial medium compared to wildtype and a *mcp1* complemented line (Figure 2A-D, Supplemental Figure S2A). While there was a lack of usage of NAD-ME-derived unlabelled pyruvate inside wildtype mitochondria, we observed a substantial and progressive increase in unlabelled citrate, 2-oxoglutarate and succinate concentrations in the extramitochondrial medium of *mpc1* samples (Figure 2E-G, Supplemental Figure S2B). Less unlabelled pyruvate was exported by *mpc1* mitochondria compared to wildtype, indicating that more NAD-ME-generated pyruvate was consumed by *mpc1* mitochondria than that by wildtype and the *mpc1* complemented line (Figure 2H). This suggests that MPC1 loss-of-function enhanced the rate of NAD-ME-derived pyruvate being oxidised by PDC significantly more than in wildtype.

**Figure 2:**
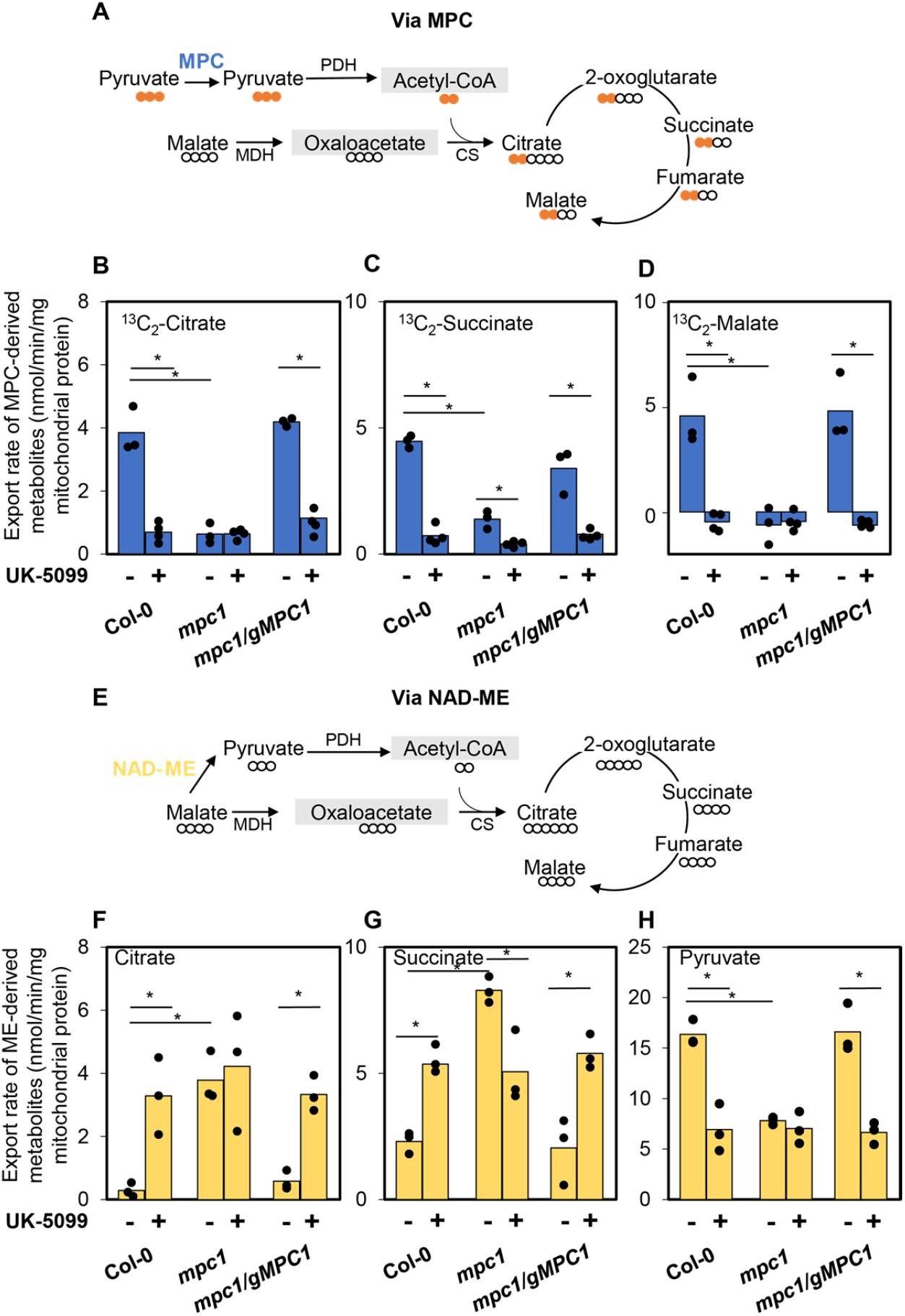
The loss of MPC1 changed the pyruvate usage pattern for generating TCA cycle intermediates. Col-0, *mpc1 and mpc1/gMPC1* mitochondria were incubated in a mixture of 500μM malate and 500μM ^13^C_3_-pyruvate at pH6.4. The isotopic incorporation patterns of labelled pyruvate and unlabelled malate into citrate via MPC (A) and via NAD-ME (E) are shown. Bar graphs show export rates of (B) ^13^C_2_-citrate (via MPC), (C) ^13^C_2_-succinate (via MPC), (D) ^13^C_2_-Malate (via MPC); (F) citrate (via NAD-ME), (G) succinate (via NAD-ME), (H) pyruvate (via NAD-ME). Quantification was carried out using SRM-MS to directly assess substrate consumption and product generation of substrate-fed mitochondria after separating mitochondria from the extra-mitochondrial space by centrifugation through a single silicon oil layer. The rates were calculated from time course values of metabolite concentration recorded in the extra-mitochondrial space after varying incubation periods. Each bar represents averaged value from three or more replicates represented by data points. Significant differences between controls and treatments are denoted by asterisks based on Student’s t-tests (*, p < 0.05).

We also included UK-5099, a non-competitive MPC inhibitor (35–38) into our feeding experiments to mimic the effect of knocking out MPC1. Figure 2 showed no significant difference in the usage of pyruvate produced from either NAD-ME or MPC between Col-0, *mpc1* and *mpc1/gMPC1* in the presence of this MPC inhibitor. All UK-5099 treated mitochondria showed a similar pattern of unlabelled pyruvate and citrate export to *mpc1* mitochondrial without UK-5099 treatment (Figure 2F, 2H, Supplemental Figure S2). UK-5099 treatment blocked transport and usage of exogenously-provided pyruvate while it enhanced the usage of pyruvate from NAD-ME in Col-0 and *mpc1/gMPC1* in the same way as knockout of MPC in *mpc1*. The flexibility in using NAD-ME-derived pyruvate to provide carbon for the TCA cycle when the primary source of pyruvate is unavailable further confirms that physical channelling is not the cause of the preferential use of imported pyruvate for supporting the TCA cycle.

To independently confirm this effect, we conducted a separate label swap mitochondrial feeding experiment with a combination of pyruvate and ^13^C_4_-malate with/without UK-5099. The results consistently showed that *mpc1* mitochondria and UK-5099 treated mitochondria were able to use pyruvate made internally to mitochondria by NAD-ME to drive the TCA cycle, but this did not occur in wildtype or the *mpc1* complemented line (Supplemental Figure S3). It was evidenced by the steady increase in NAD-ME-derived citrate and succinate concentration in the form of ^13^C_6_-citrate and ^13^C_4_-succinate (Supplemental Figure S3B). Taken together, pyruvate from NAD-ME can only be efficiently accessed by PDC when MPC1 is either not present or chemically inactivated. This means there are effectively two mitochondrial pyruvate pools in plant mitochondria that can be accessed by PDC, but it depends on the presence of pyruvate import.

### Total amount of TCA cycle metabolites produced from NAD-ME-generated pyruvate remained unchanged regardless of genotypic or treatment effect

Although the maximal activity of NAD-ME was shown to be similar in wildtype, *mpc1* and *mpc1/gMPC1* (2), we surmised the difference in unlabelled pyruvate production rate between wildtype and *mpc1* might underlie the variation in metabolic regulation under *in vitro* conditions. In order to assess NAD-ME activity of wildtype, *mpc1* and *mpc1/gMPC1* with and without UK-5099, we calculated and compared the export rates of the downstream products from NAD-ME, namely unlabelled pyruvate, citrate, succinate and 2-oxoglutarate when they were fed with 500μM ^13^C_3_-pyruvate and 500μM malate at pH 6.4. Unlabelled fumarate and malate derived from this pathway cannot be measured as malate had the same isotopically labelled form as exogenously provided malate, and fumarate production was comparatively too low to be quantifiable. Our results showed that *mpc1* and UK-5099-treated mitochondria displayed no difference in the total concentration of metabolites derived from the NAD-ME pathway (Figure 3B, Supplemental Figure S4A) while they displayed obvious deficiency in total MPC-derived metabolite amount to that of Col-0 (Figure 3A). However, the export rates of individual NAD-ME-derived metabolites were clearly different; wildtype exported the majority as NAD-ME-derived pyruvate whereas *mpc1* and UK-5099 treated mitochondria converted more than half of this pyruvate into TCA cycle metabolites (Figure 3). Swapping from supplying ^13^C_3_-pyruvate and unlabelled malate to ^13^C_4_-malate and unlabelled pyruvate showed consistent results, indicating that malic enzyme activity was similar amongst genotypes and treatments but the pyruvate generated had significantly different fates when MPC1 was absent or non-functional (Supplemental Figure S4B-C). Thus, the bias in pyruvate usage is not due to different pyruvate production rate by NAD-ME.

**Figure 3.**
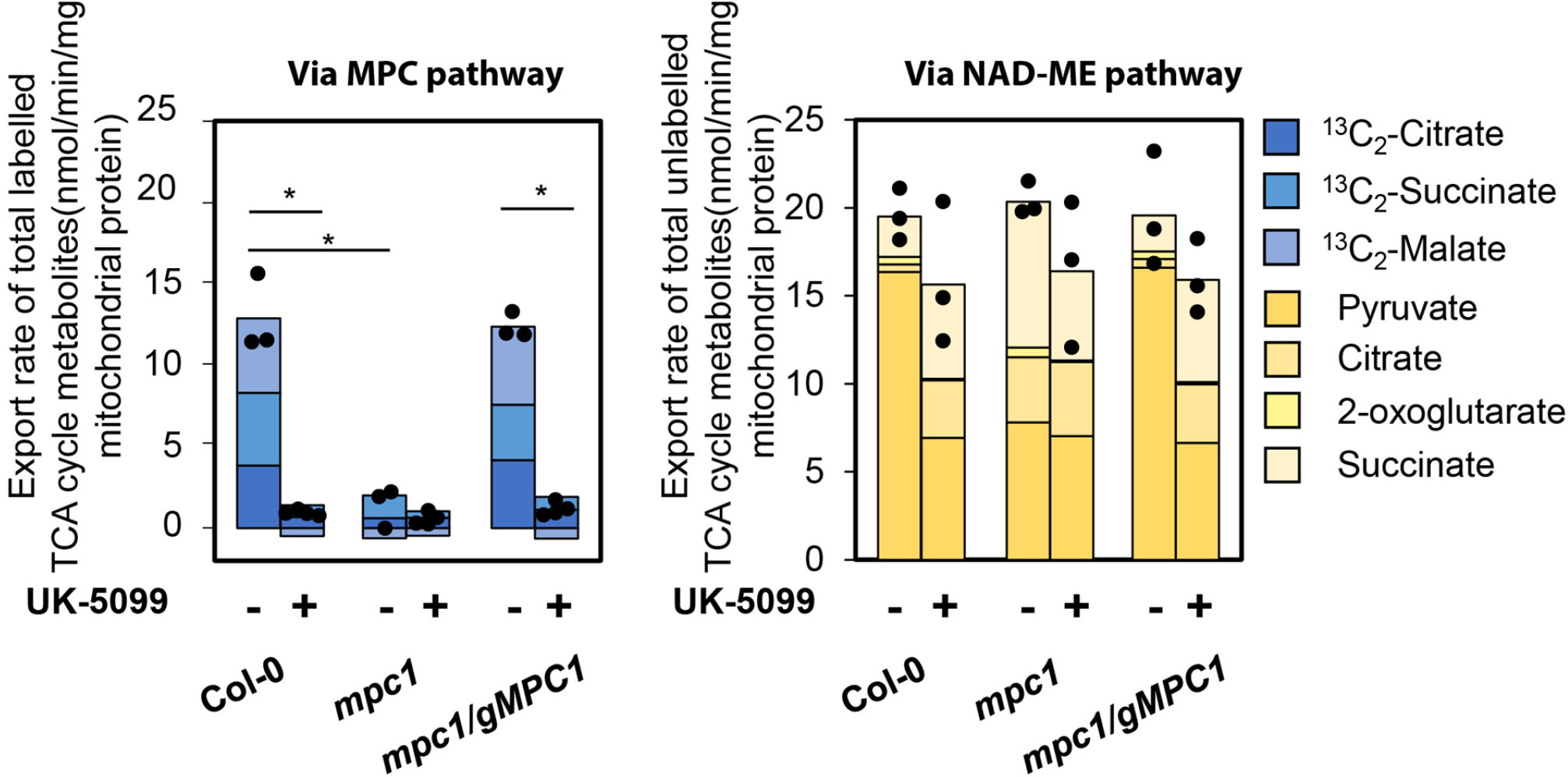
The rate of accumulation of total extra-mitochondrial NAD-ME-derived metabolites in mitochondrial feeding experiments. Bar graphs show the export rates of total extra-mitochondrial amount of MPC-derived metabolites (A) and NAD-ME-derived metabolites (B) when Col-0, *mpc1* and *mpc1*/gMPC1 mitochondria were fed with 500μM ^13^C_3_-pyruvate and 500μM malate at pH 6.4 with and without UK-5099. Quantification was carried out using SRM-MS to directly assess substrate consumption and product generation of substrate-fed mitochondria after separating mitochondria from the extra-mitochondrial space by centrifugation through a single silicon oil layer. The total amount of MPC-derived metabolites were the sum of ^13^C_2_-citrate, ^13^C_2_-succinate, ^13^C_2_-malate. The total amount of NAD-ME metabolites were the sum of unlabelled pyruvate, citrate, 2-oxoglutarate and succinate. The rates were calculated from time course values of metabolite concentration recorded in the extra-mitochondrial space after varying incubation periods. Each stacked bar represents averaged value from three or more replicates. Data points represented the total amount of metabolites exported of independent replicates. Significant differences between controls and treatments are denoted by asterisks based on Student’s t-tests (*, p < 0.05).

### Alanine aminotransferase (AlaAT) can consume NAD-ME-derived pyruvate but not MPC imported pyruvate

Our results so far suggested PDC can prioritise imported pyruvate over NAD-ME-derived pyruvate for generating TCA cycle intermediates. To explore this possibility, we introduced a competitive co-substrate, glutamate, to drive the mitochondrial alanine aminotransferase (AlaAT) in the direction of pyruvate consumption (i.e. pyruvate + glutamate --> alanine + 2-oxoglutarate) to compete for pyruvate in isolated mitochondria (Figure 4A, 4E) (39). In an equilibrium system, pyruvate imported by MPC or generated by NAD-ME would be converted by AlaAT and reduce the export rate of the corresponding citrate product. If one source of pyruvate is preferential, the export rate of citrate derived from it would be resistant to change. We found glutamate addition to mitochondria significantly deteriorated the unlabelled citrate export rate and pyruvate export rate consistent with competition for unlabelled pyruvate in Col-0 and the complemented line (Figure 4F-G, Supplemental Figure S5B). However, the rate of ^13^C_2_-citrate export,^13^C_2_-succinate and ^13^C_2_-malate did not change upon the addition of glutamate in all three genotypes, suggesting restricted access to imported pyruvate by AlaAT (Figure 4B-D, Supplemental Figure S5A). These results show that AlaAT is more likely to have access to NAD-ME-derived pyruvate than imported pyruvate in wildtype. In *mpc1* mitochondria, the export rate of unlabelled citrate derived from the NAD-ME pathway was not altered in the presence of glutamate. This suggests that, in the absence of MPC, PDC can access pyruvate from NAD-ME just with a lower rate in the presence of glutamate. Therefore, the preferential use of imported pyruvate for supporting the TCA cycle could be explained by distinct pools of pyruvate accessible to PDC depending on the rate of pyruvate import.

**Figure 4.**
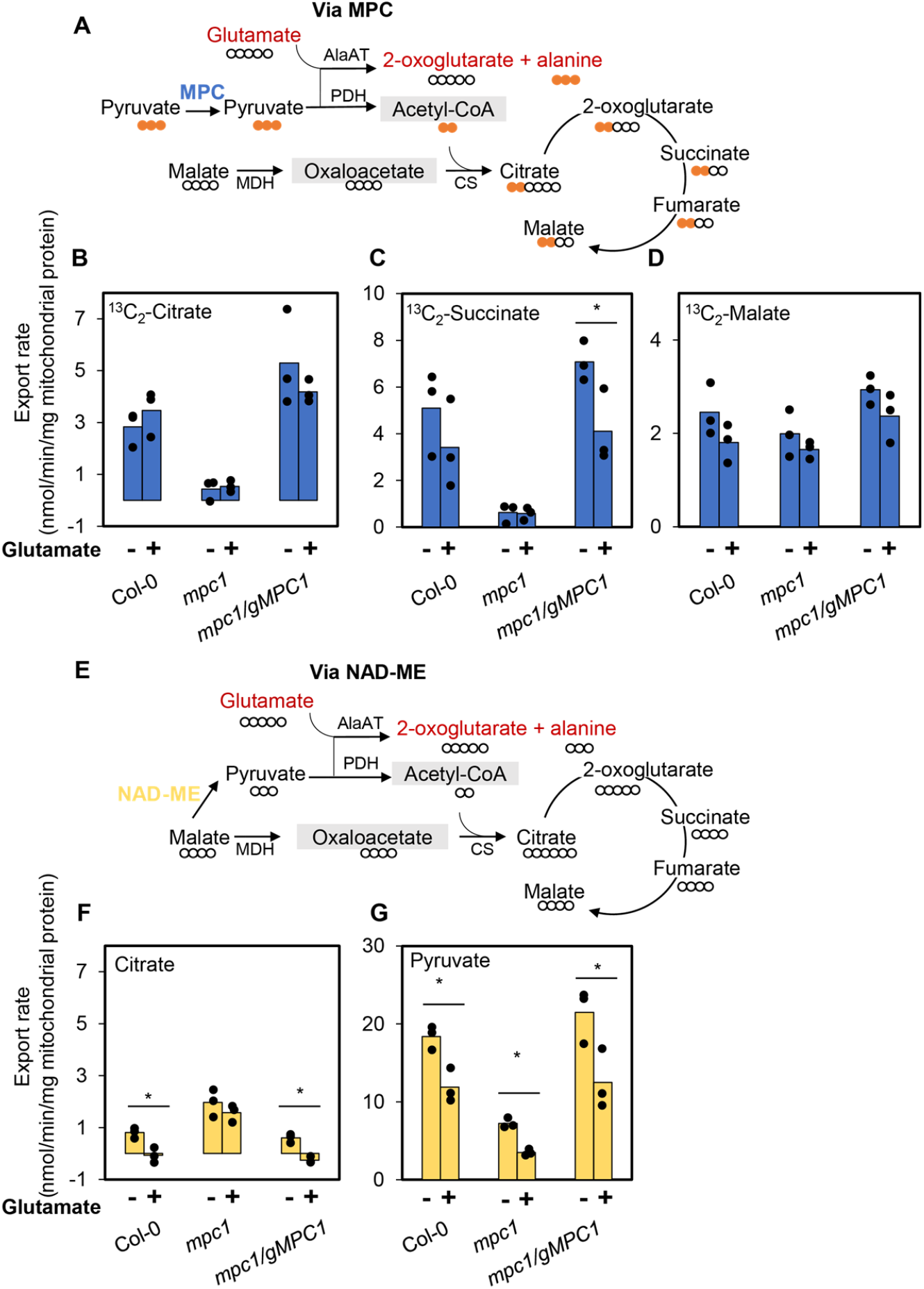
The impact of removal of pyruvate by AlaAT on citrate production in Col-0, *mpc1* and *mpc1/gMPC1* mitochondria. Mitochondria incubated with 500 μM malate and 500 μM ^13^C_3_-pyruvate with and without 500 μM glutamate at pH 6.4. The isotopic incorporation patterns of labelled pyruvate and unlabelled malate into citrate via MPC (A) and via NAD-ME (E) are shown. Bar graphs show export rates of (B) ^13^C_2_-citrate (via MPC), (C) ^13^C_2_-succinate (via MPC), (D) ^13^C_2_-malate (via MPC); (F) citrate (via NAD-ME), (G) pyruvate (via NAD-ME). Quantification was carried out using SRM-MS to directly assess substrate consumption and product generation of substrate-fed mitochondria after separating mitochondria from the extra-mitochondrial space by centrifugation through a single silicon oil layer. The rates were calculated from time course values of metabolite concentration recorded in the extra-mitochondrial space after varying incubation periods. Each bar represents averaged value from three or more replicates represented by data points. Significant differences between controls and treatments are denoted by asterisks based on Student’s t-tests (*, p < 0.05).

## Discussion

MPC plays an essential role in the mitochondrial pyruvate-supplying pathway by carrying more supply to the TCA cycle *in vivo* than AlaAT or NAD-ME (2). MPC-dependent pyruvate import alone can support normal growth and development of Arabidopsis seedlings without the presence of the other two pathways, but not AlaAT or NAD-ME alone (2) which lead to significant growth impairments. Here we show a regulatory mechanism exists to kinetically prioritise MPC over other pyruvate-supplying pathways. Specifically, we provided genetic and biochemical evidence for Arabidopsis mitochondria preferring imported pyruvate via MPC for the TCA cycle operation over pyruvate synthesised inside the mitochondrial matrix by NAD-ME. Our data indicated that imported pyruvate and matrix-derived pyruvate effectively operate as two independent pools that do not homogenously mix in the mitochondria. The rate of contribution of different pathways to the mitochondrial pyruvate pool could not explain the substantial bias of imported pyruvate usage over malate-derived pyruvate usage (Figure 1 and 4). The NAD-ME pathway provided about 90% of the total citrate in *mpc1* whereas MPC accounted for more than 90% of the total citrate in *me1*.*me2* (Supplemental Figure S6). When both pathways were available, we would expect their relative contribution to the TCA cycle to be much more evenly distributed. However, we found that 80% of citrate was still derived from MPC-transported pyruvate (Supplemental Figure S6).

### Do MPC and PDC form a metabolon?

Transient metabolons regulating the metabolic flux through the association and dissociation of components were observed for the glycolytic pathway in mammals, yeast, and plants (40–42), polyamine metabolism in plants (43), and secondary metabolic pathways in plants (44–47). Our results could be interpreted as preliminary evidence for a metabolon existing between enzymes of the mitochondrial pyruvate-supplying pathways in response to metabolic regulation and substrate availability. However, to date there is no protein-protein interaction data to support a physical association between MPC and any of the subunits of PDC, despite extensive studies being performed to define mitochondrial protein-protein interactions in plants, yeasts and mammals. These have included the use of affinity purification-mass spectrometry (AP-MS), cross-linking mass spectrometry (XL-MS), proximity-dependent biotinylation, biomolecular fluorescent complementation assay (BiFC), split luciferase, and Y2H screening assays (15, 25–32). Also, quantitative proteomics in Arabidopsis mitochondria has shown the abundance of PDC subunits are all ~5 times greater than MPC1 (48), hence at least 80% of PDC could not be associated with MPC1 at any one time. This is inconsistent with a hypothesis that MPC1 could physically associate with all PDC catalytic sites and prevent interaction with pyruvate from other sources. Rather, it appears more likely that heterogeneous zones inside mitochondria may be the reason for the observed separation of pyruvate pools. PDC has been suggested to have close association with the inner membrane as PDC isolation is more effective with detergents in animals (49, 50) and a higher portion of PDC remains associated with mitochondrial membranes in plants than other TCA cycle enzymes (51) while NAD-ME is a soluble enzyme that is free in the matrix in animals (52–54) and plants (7, 51). Additionally, immunolabelling studies in human cell cultures show that PDC is heterogeneously distributed, being found in clusters within the matrix (55).

Hypotheses that there are multiple pyruvate pools were evident in literature long before the pyruvate transporter itself was identified. U-^13^C-glucose or U-^13^C-lactate labelling in mammalian cells was used historically to develop models of compartmentation of mitochondrial metabolism that also suggests separate pyruvate pools. Most notably one originating from glucose and is used for releasable citrate, and another pool of pyruvate that seemed to function as a substrate for the TCA cycle (21). In *Saccharomyces cerevisiae*, there is also evidence that pyruvate from different origins is used for different purposes, i.e. exogenous pyruvate goes to PDH in mitochondria while glycolytic channels provide pyruvate to pyruvate decarboxylase in the cytosol (56). In *Tetrahymena*, pyruvate-derived acetyl-CoA, the product of PDC activity, is independent from acetate-derived acetyl-CoA pool (57). Pyruvate-derived CoA is oxidised for respiration, while acetate-derived CoA is used for fatty acid synthesis via citrate and fatty-acid derived acetyl-CoA is used for gluconeogenesis via malate (58). Our results show that NAD-ME-derived pyruvate and imported pyruvate do not mix in the mitochondrial matrix using an *in organello* system.

### The potential impact of pyruvate pools on primary metabolism

Metabolic pathways with separate pools of metabolic intermediates occur in all three processes of respiration; glycolysis, TCA cycle and the electron transport chain. A set of respiration-specialised metabolite pools insulated from other metabolic processes and enabling rapid and efficient energy production, is therefore a key feature of living organisms. There is good evidence that glycolytic enzymes interact with VDAC proteins of the outer mitochondrial membrane by anchoring glycolytic enzymes to the mitochondrial surface (42, 59, 60). Fructose 1,6-bisphosphate, dihydroxyacetone phosphate, and glyceraldehyde 3-phosphate are preferred to be consumed in glycolysis rather than diluted into the bulk cytosol as shown by stable isotope dilution experiments in Arabidopsis (60). Our results show that the metabolic preference for respiratory role is also true for imported pyruvate in the mitochondria. Most of pyruvate made by glycolysis readily enters the TCA cycle to immediately generate reducing power for ATP production without being used by other competing pathways (61). Similarly, in glial cells the pyruvate pool with glycolytic origin, is more closely related to mitochondrial pyruvate, which is oxidized via TCA cycle activity (62). Based on the rate of citrate export, imported pyruvate is directed with high efficiency as consistently over 80% of citrate was made from imported pyruvate (Supplemental Figure S6) to ensure the respiration efficiency.

An important component of metabolic regulation is specialization. Our results show that in plants, the imported pyruvate pool is designated to provide a carbon backbone to make citrate, whereas NAD-ME-derived pyruvate is destined to be exported to the cytosol for other cellular roles (Figure 5). The activity of NAD-ME and the mitochondrial pyruvate exporter ensures the integrity of the photosynthetic metabolism in C4 plants by the recycling of carbon intermediates. The export of NAD-ME-derived pyruvate from mitochondria is essential for PEP synthesis in the chloroplasts to accept CO_2_ in the mesophyll cells (63–66). NAD-ME releases CO_2_ from malate which allow carbon incorporation by Rubisco in the bundle sheath cells. The whole process helps to minimize photorespiration and energy wastage, thereby increasing plant yield. While the identity of the plant mitochondrial pyruvate exporter is currently unknown, our results suggest that the metabolic arrangement of pyruvate pools already operate in C3 plants to facilitate the non-mitochondrial usage of NAD-ME-derived pyruvate (67–69), albeit at a much lower rate than in C4 plants (70). In C3 plants like rice and Arabidopsis, pyruvate generated by NAD-ME from imported malate is exported to be recycled into phosphoenolpyruvate in the cytosol and plastids to restore the pH balance in response to water stress (71, 72). Moreover, pyruvate exported from mitochondria is potentially the main source of acetyl CoA to synthesise fatty acids in plastids for generating essential cellular components and signals (73–75). In summary, our data shows a process by which there is a specialized role of NAD-ME-derived pyruvate, when MPC is operating, not to supply substrates for respiration but to maintain photosynthesis, biosynthesis and potentially mediate metabolic stress responses. This metabolic specialisation explains the contribution of NAD-ME to reduction of the matrix NADH pool and thus some respiratory flux in Arabidopsis and other plants (6–8) but argues that the assumption of NAD-ME being a main source of pyruvate for the TCA-cycle is not generally applicable.

**Figure 5.**
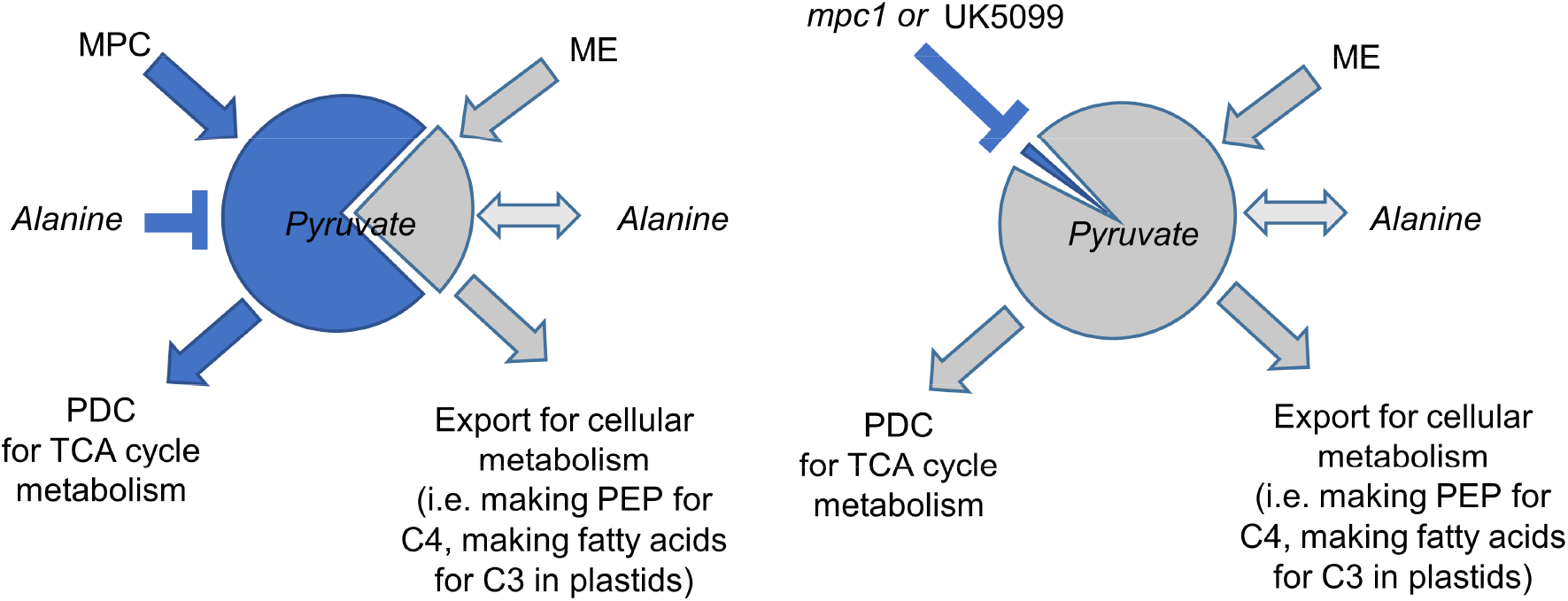
Schematic presentation of the source and sink of mitochondria pyruvate pools via MPC (Blue) and NAD-ME (Grey). In wildtype plants (left), the imported pyruvate pool is prioritised to be accessed by PDC to make acetyl-CoA which is then condensed with OAA to enter the TCA cycle while most of the NAD-ME-derived pyruvate pool is exported from the mitochondria for other cellular processes. In *mpc1* (or UK5099 treated wildtype plants) the imported pool is no longer available and the NAD-ME-derived pyruvate pool is used for both TCA cycle metabolism and export for other purposes. AlaAT can readily use the NAD-ME-derived pyruvate pool to make alanine in both wildtype and MPC-deficient mitochondria, but cannot access pyruvate imported by MPC.

Our findings show that pyruvate pools are both separated but also metabolically flexible, proven by genetics and inhibitors. This metabolic distinction allows individual turn over, flux regulation, equilibrium and hence specialised response of each metabolic pool to cellular and environmental stimuli. The presence of distinct mitochondrial pyruvate pools due to mitochondrial anatomical and biochemical features suggests similar regulation of other mitochondrial metabolites could exist in order to improve the efficiency and dynamic nature of the respiratory pathway (76). It has been suggested previously that there could be two different malate pools in plant mitochondria when castor bean mitochondria were incubated with ^14^C-pyruvate and malate. TCA cycle-generated ^14^C-malate is exported as no appreciable radiolabel can be found in CO_2_ (77). The existence of a MDH-CS metabolon also indicates that there could be at least two pools of malate, one generates OAA for the TCA cycle and the other consumed by other pathways such as NAD-ME (12–15). There is also a prima-facie case for at least two pools of citrate due to a citrate synthase–citrate exporter interaction (78). Metabolic plasticity is not compromised by having two mitochondrial pyruvate pools. By having a flexible secondary pyruvate pool, plant mitochondria have the ability to use NAD-ME-derived pyruvate to generate TCA intermediates as a fine-tuning regulation mechanism of pyruvate metabolism rather than being alternative substrates for respiration. Understanding the conditions and mechanism that enable NAD-ME contribution to the TCA cycle metabolism *in vivo* will be beneficial for the incorporation of the superior C4 characteristics into future C3 crops.

## Materials and Methods

### Plant material and growth conditions

The MPC1 T-DNA insertion line SALK008465 was obtained from the Arabidopsis Biological Resource Center (https://abrc.osu.edu/). *mpc1* and *me1*.*me2* seeds were previously characterised and published (2, 7) and *me1*.*me2* seeds were obtained from Professor Verónica G. Maurino (University of Bonn). *me1*.*me2*.*mpc1* and *mpc1/gMPC1* were generated and confirmed as described previously (2).

Arabidopsis seeds were surface sterilized and dispensed into one-half strength Murashige and Skoog liquid media (1.1g/L agar, 0.4 g/L MES, 10g/L sucrose) within enclosed, sterilized 100 mL polypropylene containers. The containers were rotated on the shaker in the long-day conditions with 16 hours light, 8 hours dark and 60% humidity (110 μmol s−1 m−1 light intensity with tubular fluorescent lighting) and seedlings were harvested after two weeks.

### Isolation of mitochondria

Mitochondria were isolated from 2-week-old Arabidopsis seedlings as described previously (79).

### Substrate feeding of isolated mitochondria

The detailed methods and materials for MS-based mitochondria feeding assays are described previously (80). In short, 100 μg isolated mitochondria were mixed with substrates (a mixture of pyruvate and malate), cofactors (2mM NAD^+^, 0.2 mM TPP and 0.012 mM CoA) and 1mM ADP (for ATP synthesis) in a final volume of 200 μl. At specified time, this reaction mixture was layered on top of silicon oil (AR200, 100 μl) which was layered above the stopping sucrose solution (0.5 M sucrose, pH 1.0). Substrate transport was stopped by rapid centrifugation (12 000 *g* for 3 min) to harvest the mitochondria at the bottom of the tube. 5 μl of the extra-mitochondrial medium (the top layer) was collected and extracted for quantitative analysis by LC-SRM-MS.

### Analyses of metabolites by LC-SRM-MS

Samples were analysed by an Agilent 1100 HPLC system coupled to an Agilent 6430 Triple Quadrupole (QQQ) mass spectrometer equipped with an electrospray ion source as described previously (2). Chromatographic separation was performed on a Kinetex C18 column, using 0.1% formic acid in water (solvent A) and 0.1% formic acid in methanol (solvent B) as the mobile phase for binary gradient elution. The elution gradient was 18% B at 1 min, 90% B at 10 min, 100% B at 11 min, 100% B at 12 min, 18% B at 13 min, and 18% B at 20 min. The column flow rate was 0.3 mL/min; the column temperature is 40 °C, and the autosampler was kept at 10 °C. Selective reaction monitoring (SRM) transitions for targeted TCA cycle metabolites and their isotopically labelled versions are shown previously (2). Data acquisition was performed using Agilent MassHunter Workstation Data Acquisition software. Metabolite quantitation of both unlabelled and labelled metabolites was carried out based on calibration curves obtained with unlabelled authentic standards and normalized against internal standards.

### Interactome analyses

The improved Y2H high-throughput binary interactome mapping liquid pipeline described (81) is an adaptation of a previously developed interactome (82). The same low copy number yeast expression vectors expressing DB-X and AD-Y hybrid proteins and the two yeast two hybrid strains, *Saccharomyces cerevisiae* Y8930 and Y8800 were used. The reporter genes GAL2-ADE2 and LYS2::GAL1-HIS3 are integrated into the yeast genome. Expression of the GAL1-HIS3 reporter gene was tested with 1 mM 3AT (3-amino-1,2,4-triazole, a competitive inhibitor of the HIS3 gene product). Y8800 MATa and Y8930 MATα yeast strains were transformed with AD-Y and DB-X constructs, respectively and DB-X strains were tested for to be auto-activation of the GAL1-HIS3 reporter gene in the absence of AD-Y plasmid. MPC1 was cloned into DB-X construct acting as baits and screened against 12000 proteins in Arabidopsis library cloned into AD-Y construct prior to Y2H screening.

Briefly, DB-X baits expressing yeasts were individually grown (30°C for 72 h) into 50-mL polypropylene conical tubes containing 5 mL of fresh selective media (Sc-Leucine; Sc-Leu), then pooled (max 50 individual bait yeast strains) and 50 μL plated into 384-well low profile microplates. Glycerol stocks of the (AD)-AtORFeome collection corresponding to 127 96-well plates were thawed, replicated using the colony picker Qpix2 XT into 32 384-well plates filled with 50 μL of fresh selective media (Sc-Tryptophane; Sc-Trp) and incubated at 30 °C for 72 h. Culture plates corresponding to the DB-baits pools and AD-collection were replicated into mating plates filled with YEPD media and incubated at 30 °C for 24 h. Mating plates were then replicated into screening plates filled with 50 μL of fresh Sc-Leu-Trp-Histidine + 1 mM 3AT media and incubated at 30 °C for 5 days. Only diploid yeast with interacting couples can growth in this media. In order to identify primary positives, the OD600 of the 384-well screening plates was measured using a microplate-reader Tecan Infinite M200 PRO. Yeast cultures identified as positive interactions were picked from selective media and protein pairs were identified by de-pooling of DB-baits in a targeted matricial liquid assay in which all the DB-baits were individually tested against all the positive AD-proteins. Identified pairs were cherry-picked and checked by DNA sequencing.

### Statistical analysis

All statistical analyses were performed using the two-sided *t* test function built in Excel 2010. Statistical tests and the number of biological replicates are indicated in figure legends. Biological replicates indicate samples that were collected from different batches of plants grown under the same conditions except biological replicates for transcript analysis and metabolite analysis were samples collected from different plants grown at the same time.

## Acknowledgements

This work is supported by the Australian Research Council Centre of Excellence in Plant Energy Biology (CE140100008) and X.H.L. is a Forrest Scholar supported by the Forrest Research Foundation and a receiver of Research Training Program scholarships from the Department of Education, Skills and Employment in the Australian Government.

## Figures

**Supplemental Figure S1.**
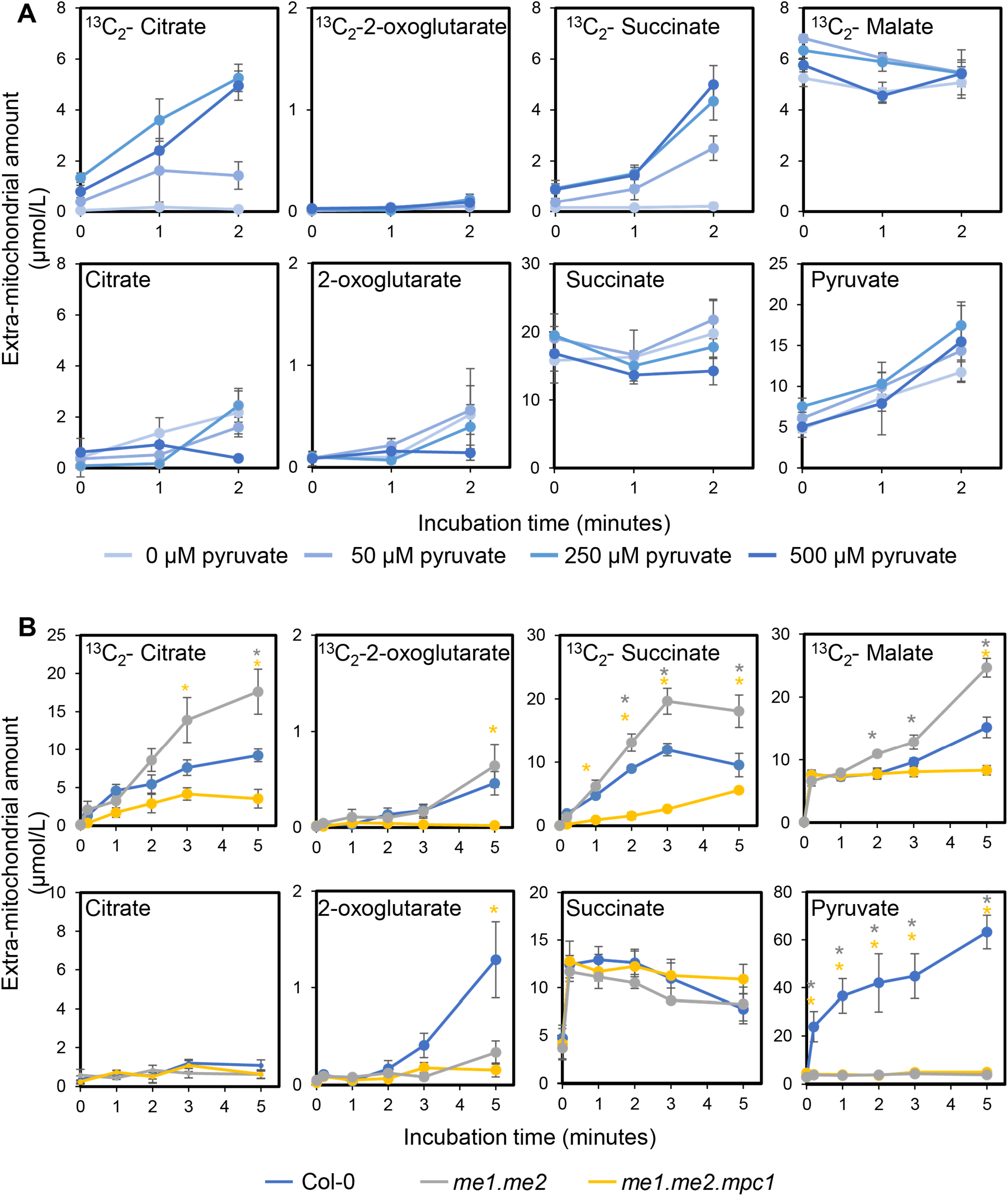
^13^C_3_-Pyruvate and malate feeding to isolated mitochondria of Col-0, *me1*.*me2* and *mpc1*.*me1*.*me2*. Time courses of metabolite concentrations in the extra-mitochondrial space of isolated mitochondria incubated with 500 μM ^13^C_3_-pyruvate and 500 μM malate via the MPC pathway (A) and via the NAD-ME pathway (B). All experiments were conducted in the presence of ADP at pH 6.4 to initiate substrate uptake and consumption by both pathways. Metabolic reaction was stopped by centrifugation through a single silicon oil layer by which the mitochondrial pellet was separated from the extra-mitochondrial medium. Unused substrate and exported products in the extra-mitochondrial medium were quantified using LC-SRM-MS. Each data point represents averaged value from three or more biological replicates with error bars indicate standard error (n≥3). Significant differences between *mpc1*, Col-0 and *mpc1/gMPC1* are denoted by asterisks based on Student’s t-tests (*, p < 0.05). (Supports Figure 1).

**Supplemental Figure S2.**
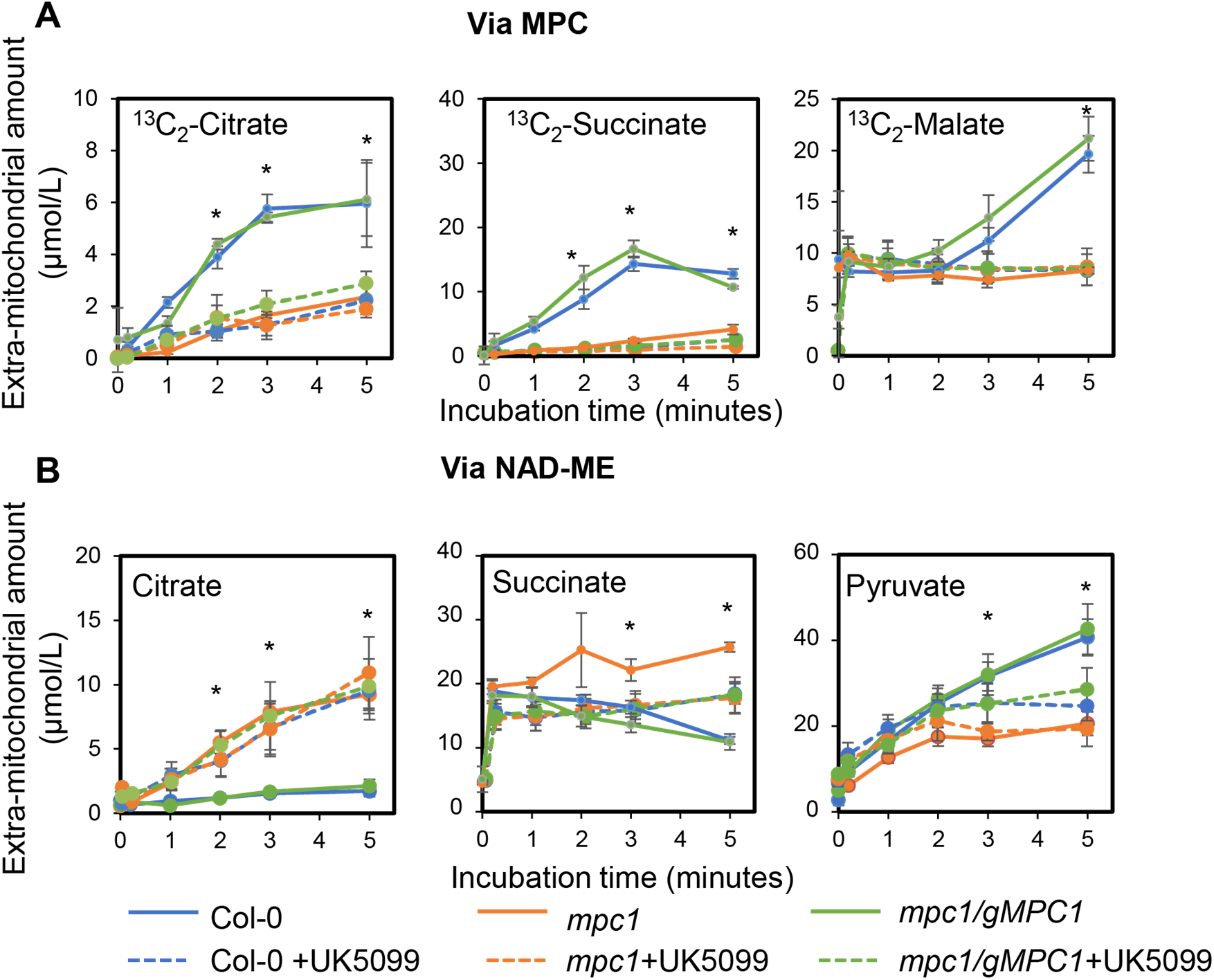
^13^C_3_-Pyruvate and malate feeding to isolated mitochondria of Col-0, *mpc1* and *mpc1/gMPC1*. Time courses of metabolite concentrations in the extra-mitochondrial space of isolated mitochondria incubated with 500 μM ^13^C_3_-pyruvate and 500 μM malate via MPC pathway (A) and via NAD-ME pathway (B). All experiments were conducted in the presence of ADP at pH 6.4 to initiate substrate uptake and consumption by both pathways. Metabolic reaction was stopped by centrifugation through a single silicon oil layer in which the mitochondrial pellet was separated from the extra-mitochondrial medium. Unused substrate and exported products in the extra-mitochondrial medium were quantified using LC-SRM-MS. Each data point represents averaged value from three or more biological replicates with error bars indicate standard error (n≥3). Significant differences between *mpc1*, Col-0 and *mpc1/gMPC1* are denoted by asterisks based on Student’s t-tests (*, p < 0.05) (Supports Figure 2).

**Supplemental Figure S3.**
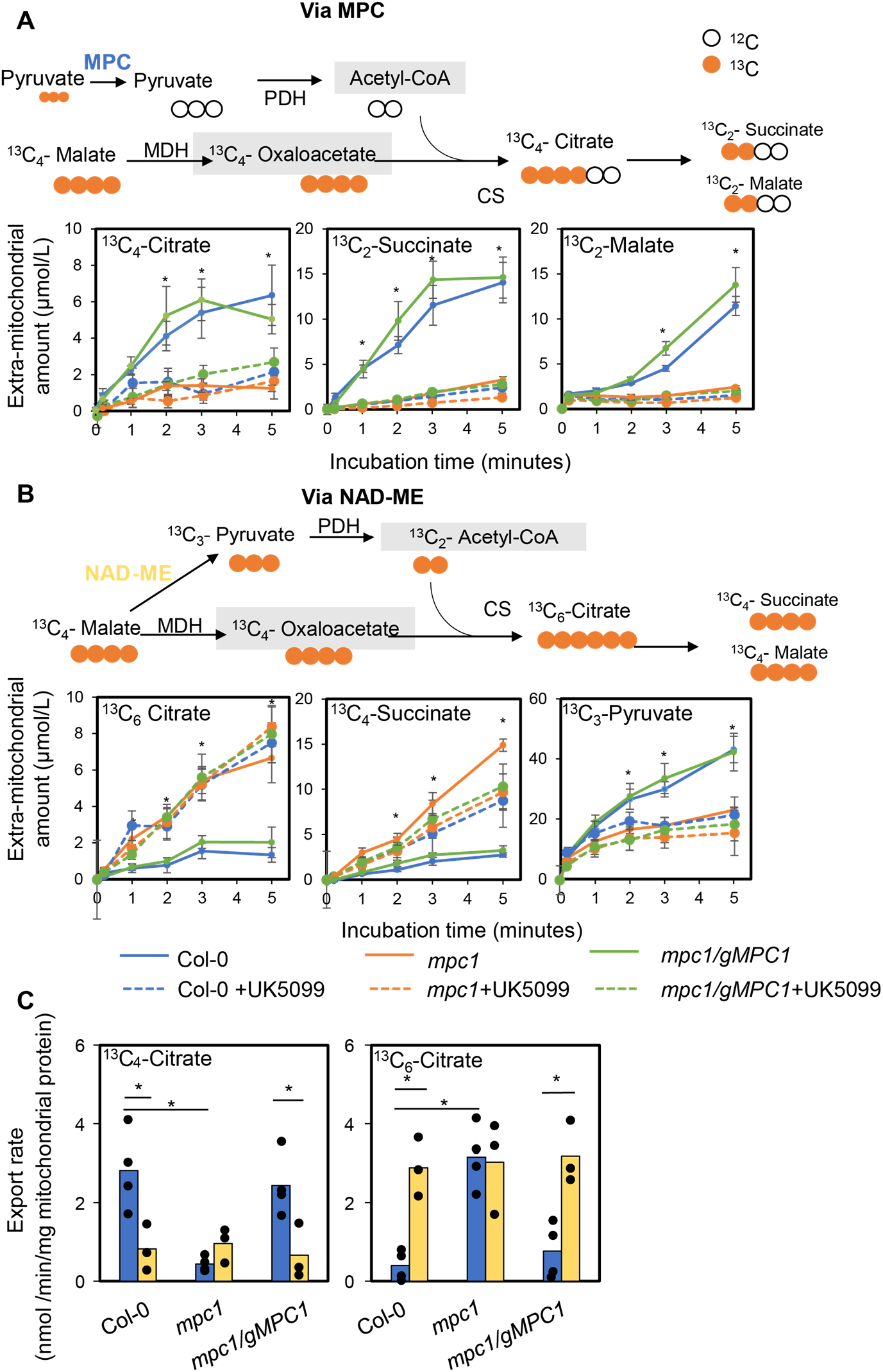
Pyruvate and ^13^C_4_-malate feeding to isolated mitochondria of Col-0, *mpc1* and *mpc1/gMPC1*. Time courses of metabolite concentrations in the extra-mitochondrial space of isolated mitochondria incubated with 500 μM pyruvate and 500 μM ^13^C_4_-via MPC pathway (A) and via NAD-ME pathway (B). All experiments were conducted in the presence of ADP at pH 6.4 to initiate substrate uptake and consumption by both pathways. Metabolic reaction was stopped by centrifugation through a single silicon oil layer in which the mitochondrial pellet was separated from the extra-mitochondrial medium. Unused substrates and exported products in the extra-mitochondrial medium were quantified using LC-SRM-MS. Each data point represents averaged value from three or more biological replicates with error bars indicate standard error (n≥3). Significant differences between *mpc1*, Col-0 and *mpc1/gMPC1* are denoted by asterisks based on Student’s t-tests (*, p < 0.05) (C) Bar graphs show the rates calculated from time course values of metabolite concentration recorded in the extra-mitochondrial space after varying incubation periods. Each bar represents averaged value from three or more replicates represented by data points. Significant differences between controls and treatments are denoted by asterisks based on Student’s t-tests (*, p < 0.05). (Supports Figure 2).

**Supplemental Figure S4.**
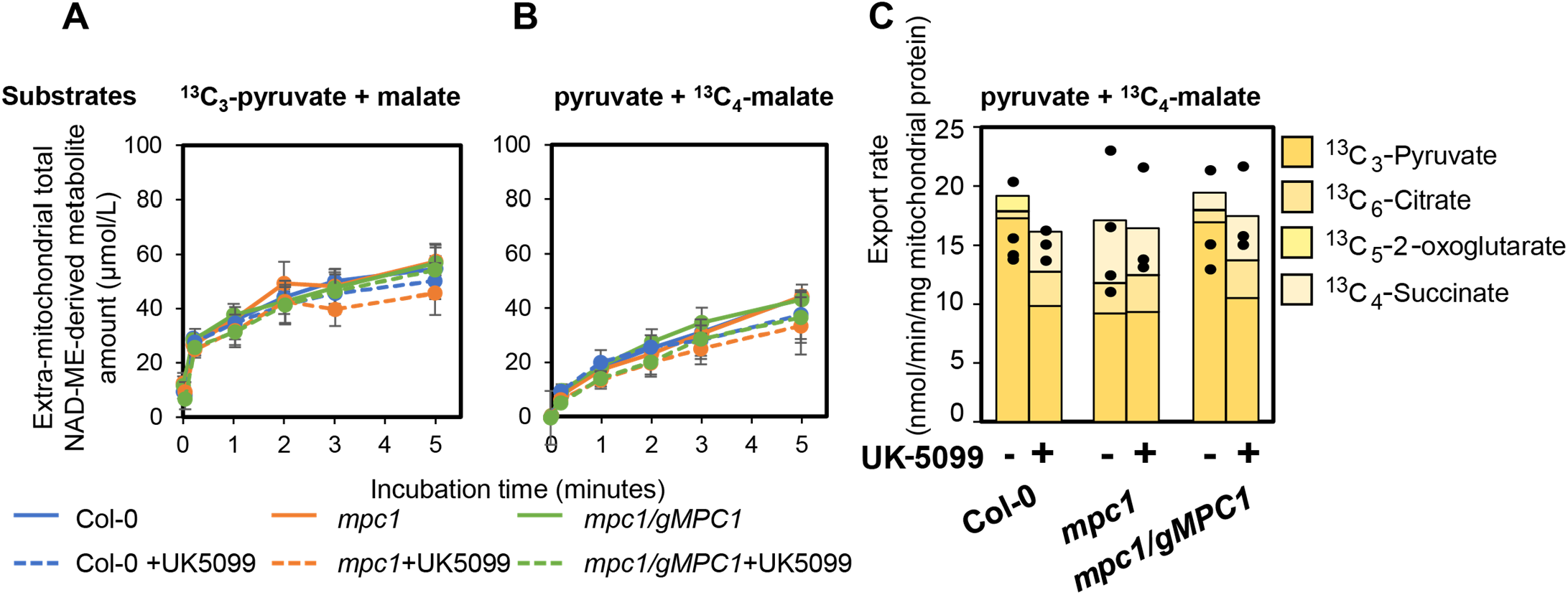
The total amount and the rate of metabolites exported from mitochondria that were made via the NAD-ME pathway. The total amount of NAD-ME derived metabolites was calculated from time course experiments of either pyruvate and ^13^C_4_-malate feeding (A, including unlabeled citrate, 2-oxoglutarate, succinate, pyruvate) or pyruvate and ^13^C_4_-malate feeding (B, including ^13^C_6_-citrate, ^13^C_5_-2-oxoglutarate, ^13^C_4_-succinate, ^13^C_3_-pyruvate) to isolated mitochondria of Col-0, *mpc1* and *mpc1/gMPC1*. All experiments were performedin the presence of ADP at pH 6.4 to initiate substrate uptake and consumption by both pathways. Metabolic reaction was stopped by centrifugation through a single silicon oil layer in which the mitochondrial pellet was separated from the extra-mitochondrial medium. Unused substrates and exported products in the extra-mitochondrial medium were quantified using LC-SRM-MS. Each data point represents averaged value from three or more biological replicates with error bars indicate standard error (n≥3). Significant differences between *mpc1*, Col-0 and *mpc1/gMPC1* are denoted by asterisks based on Student’s t-tests (*, p < 0.05). (C) Bar graphs show the calculated export rate of all metabolites combined which were made from ME-derived pyruvate after 5 minutes feeding the mitochondria with pyruvate and ^13^C_4_-malate. Each stacked bar represents averaged value of the indicated metabolite from three or more replicates. Data points represented the total amount of ME-derived metabolites exported in independent replicates. Significant differences between controls and treatments are denoted by asterisks based on Student’s t-tests (*, p < 0.05). (Supports Figure 3).

**Supplemental Figure S5.**
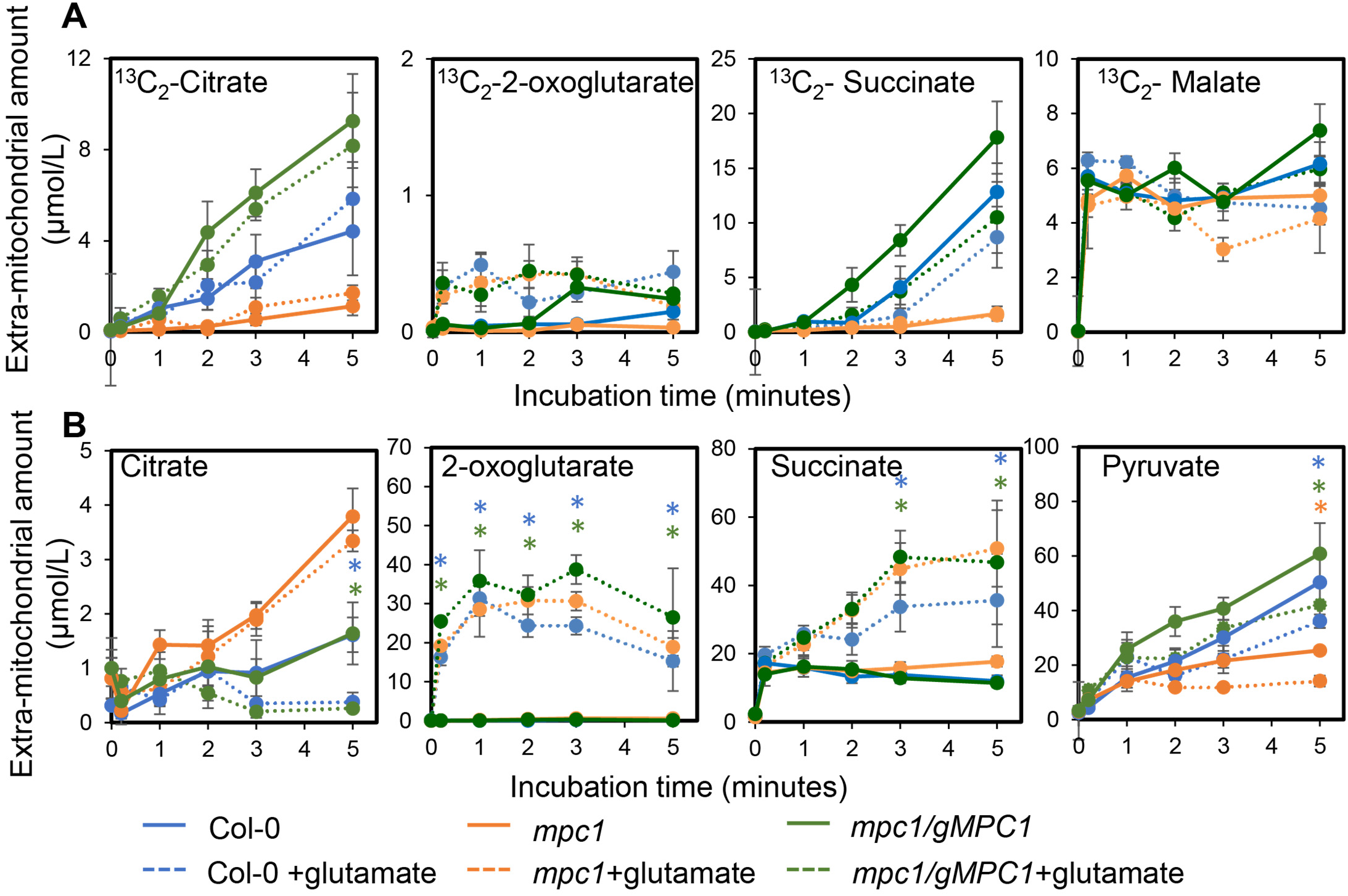
^13^C_3_-Pyruvate and malate feeding to isolated mitochondria of Col-0, *mpc1* and *mpc1/gMPC1* with or without the addition of glutamate. Time courses of metabolite concentrations in the extra-mitochondrial space of isolated mitochondria incubated with 500 μM ^13^C_3_-pyruvate and 500 μM malate with or without the addition of glutamate. All experiments were conducted in the presence of ADP at pH 6.4 to initiate substrate uptake and consumption via both MPC and NAD-ME pathways. Metabolic reaction was stopped by centrifugation through a single silicon oil layer in which the mitochondrial pellet was separated from the extra-mitochondrial medium. Unused substrate and exported products in the extra-mitochondrial medium were quantified using LC-SRM-MS. Line graphs show the amount of ^13^C_2_-citrate (A), citrate (B) and pyruvate (C) during 5-minute incubation. Each data point represents averaged value from three or more biological replicates with error bars indicate standard error (n≥3). Significant differences between controls (straight lines) and treatments (dotted lines) are denoted by asterisks based on Student’s t-tests (*, p < 0.05) (Supports Figure 4).

**Supplemental Figure S6.**
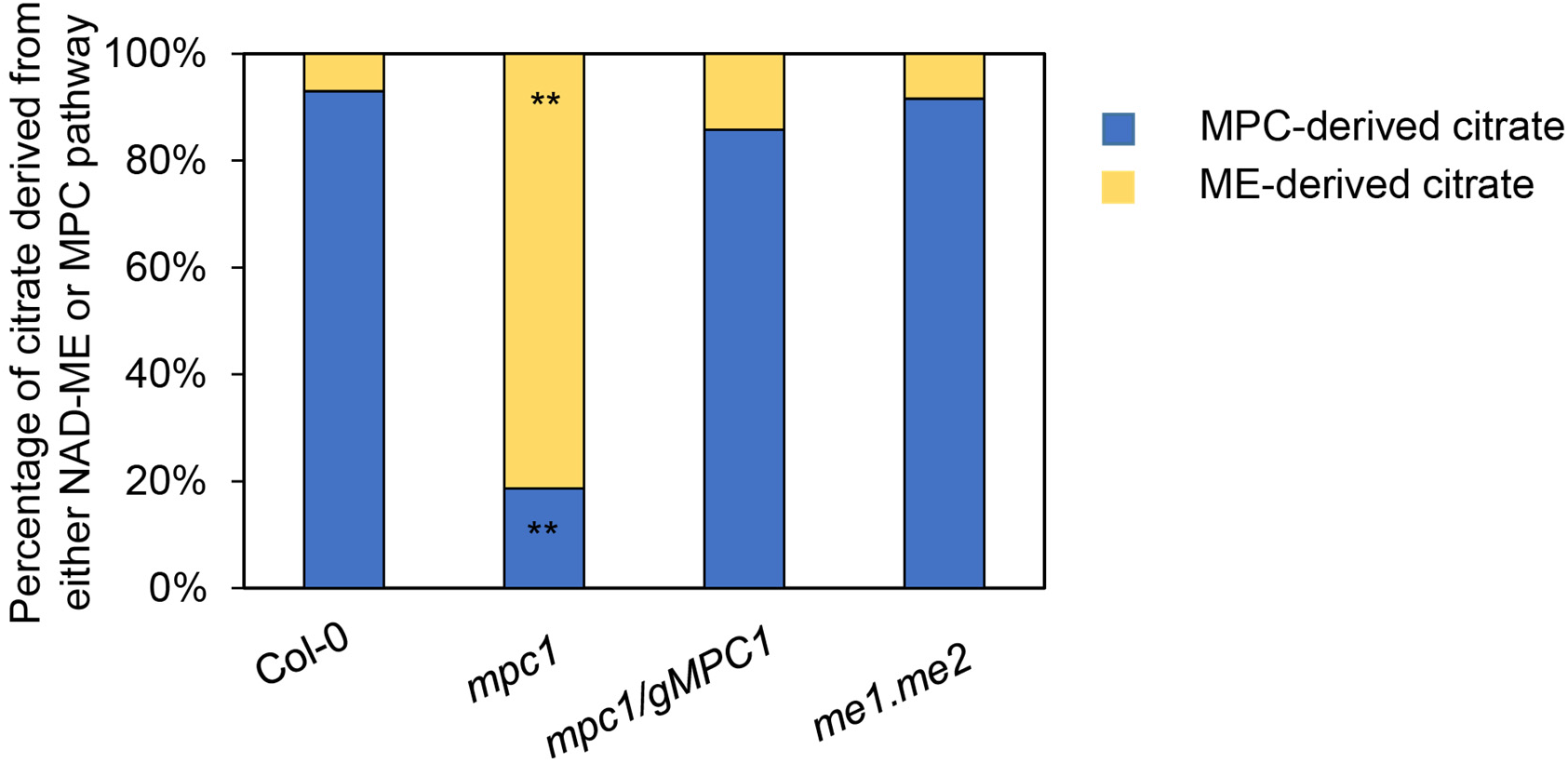
The proportion of citrate derived from difference sources of pyruvate. Isolated mitochondria incubated with 500 μM ^13^C_3_-pyruvate and 500 μM malate with or without the addition of glutamate in the presence of ADP at pH 6.4 to initiate substrate uptake and consumption to initiate substrate uptake and consumption via MPC and NAD-ME pathways. Metabolic reaction was stopped by centrifugation through a single silicon oil layer by which the mitochondrial pellet was separated from the extra-mitochondrial medium. The amount of citrate (labelled and unlabelled) in the extra-mitochondrial medium were quantified using LC-SRM-MS. Bar graphs show the percentage of ^13^C_2_-citrate (blue) and citrate (yellow) derived from the MPC and NAD-ME pathways, respectively. Each bar is the averaged value from three or more biological replicates (n≥3). Significant differences between wildtype and mutants are denoted by asterisks based on Student’s t-tests (*, p < 0.05) (Supports Figure 5).

## Notes

**Competing Interest Statement:** There is no conflict of interests

### Competing Interest Statement

The authors have declared no competing interest.

